# An Interpretable EEG Machine Learning Prototype for Alzheimer’s Disease Classification

**DOI:** 10.64898/2026.09.12.751104

**Authors:** Atrim Das, Anamika Chakravarty

## Abstract

Resting-state electroencephalography (EEG) can capture the slowing of neural oscillations associated with Alzheimer’s disease (AD), but many machine-learning studies remain difficult to inspect, reproduce, or test. This study developed an interpretable, subject-level AD versus healthy-control classifier from the dataset OpenNeuro ds004504 and deployed it as a public research prototype. Preprocessed eyes-closed EEG recordings from 36 people with AD and 29 healthy controls were analysed. Welch power spectral density estimates were used to generate absolute and relative bandpower summaries, theta/alpha and delta/alpha ratios, a slow/fast ratio, and signal-level descriptors. Five-fold stratified cross-validation compared two Random Forest configurations, EEG Logistic Regression, an age-only baseline, and an EEG-plus-age model. EEG Logistic Regression gave the strongest mean fold-wise performance: accuracy 0.846, balanced accuracy 0.848, F1 score 0.849, and ROC-AUC 0.948. Age alone was near chance, while adding age did not improve the EEG-only model. The feature pattern was consistent with AD-related EEG slowing, including higher slow-wave-related ratios and lower relative alpha power. The trained pipeline was deployed as a Gradio application on Hugging Face Spaces. The result is a reproducible research and educational prototype, not a clinical diagnostic device, and requires external validation before any clinical interpretation.

## Introduction

Alzheimer’s disease is not diagnosed from a single signal. Clinical assessment usually brings together history, cognitive testing, neurological examination, and, where available, imaging or biomarker evidence. That complexity is appropriate. It also means that many diagnostic pathways depend on specialist services, time, and unevenly available resources. Resting-state electroencephalography offers a different kind of information. It records cortical electrical activity non-invasively and can be acquired with routine clinical EEG systems. Its value in Alzheimer’s disease is not that it replaces established diagnostic work-up, but that it provides a low-cost physiological measure that can be analysed quantitatively. A recurring finding in the Alzheimer’s EEG literature is a shift toward slower oscillatory activity. Relative delta and theta activity often increase, while alpha activity is reduced or displaced. These group-level effects are not uniform in every study, and they are influenced by disease stage, recording protocol, medication, artefact handling, and feature definition. Even so, the broad pattern has been reported often enough to make spectral analysis a reasonable starting point for transparent machine-learning work [1,2]. The practical attraction is straightforward: the same spectral measures that underpin a classifier can also be shown to a reader, discussed against known physiology, and checked for implausible directions. The present study used OpenNeuro ds004504, a public dataset described by Miltiadous et al. [3]. It contains resting-state, eyes-closed EEG recordings from people with Alzheimer’s disease, frontotemporal dementia, and cognitively normal controls. The dataset is useful for secondary analysis because its files are BIDS-organised, participant metadata are supplied, and preprocessed derivatives are available. The source article also reported benchmark classification results based on relative bandpower features. This project was built around that gap. The aim was not to claim a new clinical diagnostic system or to compete directly with deep-learning studies using a different unit of analysis. The aim was to construct a clear pathway from public EEG data to an interpretable, reproducible, and accessible research prototype. The analysis retained one feature row per participant rather than treating short EEG windows from the same recording as independent observations. It extracted features that have a simple physiological interpretation, compared flexible and constrained classifiers, tested age as a competing explanation, and deployed the final model in a web application. Four features distinguish the present work from a basic reproduction of the source dataset paper. First, the analysis used a subject-level feature table, which makes the cross-validation unit explicit and avoids inflating the sample through overlapping EEG segments. Second, it compared an original Random Forest with a deliberately more conservative forest and a regularised Logistic Regression model. Third, it included an age-only baseline and an EEG-plus-age comparison. These controls matter because age and diagnosis can be entangled in neurodegeneration datasets. Fourth, the trained model was packaged into a public Gradio application hosted on Hugging Face Spaces. The app does not diagnose Alzheimer’s disease, but the primary objective was to classify Alzheimer’s disease versus healthy-control EEG profiles using interpretable spectral and oscillatory features. Secondary objectives were to test age as a competing explanation and to make the workflow accessible as a public research demonstration.

## Methods

### Dataset and analytical cohort

This was a secondary analysis of public, de-identified EEG data from OpenNeuro ds004504. The source cohort included 88 participants: 36 with Alzheimer’s disease, 23 with frontotemporal dementia, and 29 healthy controls [3]. The present analysis used a binary Alzheimer’s disease versus healthy-control subset. The frontotemporal dementia group was excluded because the deployed prototype was designed for a two-class demonstration and had not been trained or evaluated as a three-class dementia model. The resulting analytical cohort contained 65 participants: 36 with Alzheimer’s disease and 29 healthy controls. The recordings were resting-state, eyes-closed scalp EEG acquired using a 19-channel international 10–20 montage at 500 Hz. Dataset metadata included age, gender, diagnostic group, and Mini-Mental State Examination score. No new participant recruitment, clinical assessment, or data collection occurred in this project. The analysis used the preprocessed EEGLAB derivative .set files rather than the raw recordings. This was a deliberate decision. According to the dataset descriptor, the derivative workflow had already included band-pass filtering, mastoid-based re-referencing, artefact subspace reconstruction, independent component analysis, and automated identification and removal of ocular and jaw-related components [3]. Using these derivatives reduced the risk of presenting a lightly processed raw dataset as if it had undergone a complete denoising protocol within this project.

### Ethics statement

This study involved secondary analysis of publicly available, de-identified data from the OpenNeuro dataset ds004504. No participants were recruited, and no new human-subject data were collected for the present study. Ethical approval and informed-consent procedures governing the original data collection are described in the source dataset publication [3].

### Secondary standardisation and quality control

The derivative files were loaded in MNE-Python [4]. A representative recording was checked for the expected 19 channels, 500 Hz sampling frequency, and a plausible 1 to 45 Hz power spectrum. This was a quality-control check for file consistency rather than an exclusion procedure. A secondary standardisation step was then applied to each derivative recording. EEG channels were retained, the recording was re-referenced to the average reference, and a 1 to 45 Hz band-pass filter was applied. This step was intentionally limited. It should not be described as a replacement for the source authors’ artefact handling. Its role was to ensure that every file entered the feature-extraction function with the same channel selection, reference convention, and frequency range.

### Spectral and oscillatory feature engineering

Power spectral density was estimated with Welch’s method, which averages periodograms from shorter segments to obtain a more stable frequency-domain estimate [5]. The implementation used two-second segments. For each participant, power was calculated across five conventional bands: delta (1 to 4 Hz), theta (4 to 8 Hz), alpha (8 to 13 Hz), beta (13 to 30 Hz), and gamma (30 to 45 Hz). The selected ranges differ slightly from the source dataset paper, which used a 0.5 Hz lower bound and a beta/gamma boundary at 25 Hz. The present ranges were chosen to align with the secondary 1 to 45 Hz filter and to keep the feature definition simple and reproducible. For every band, values were summarised using mean, standard deviation, median, and maximum absolute power. Relative bandpower was added to reduce dependence on overall signal amplitude and recording gain. Three ratio measures were included because they directly reflect the slow-wave shift often discussed in Alzheimer’s EEG studies: the theta/alpha ratio, the delta/alpha ratio, and the slow/fast ratio. The first two compare slow activity with alpha, which is commonly reduced in Alzheimer’s disease. The slow/fast ratio summarises the balance between lower-frequency and faster-frequency components. The feature table also retained signal-level mean, standard deviation, variance, and peak-to-peak values. These were included as broad descriptors of the signal and were not assumed to be disease-specific markers. The extraction procedure produced 38 columns, including the participant identifier. After joining features to metadata and retaining the two diagnostic groups, the model matrix contained 65 participants and 37 EEG-derived predictor features. Figure 1 places the extracted features in context. Panel A is a schematic adaptation of group-level spectral patterns reported by Miltiadous et al. [3] and is included to illustrate the physiological rationale for selecting slow-wave and alpha-related features. Panel B presents descriptive Alzheimer’s disease minus healthy-control mean differences for selected native-scale features. These values were used to visualise the direction of model-relevant feature variation and were not intended as formal inferential evidence of between-group biomarker differences. No univariate hypothesis testing, effect-size estimation, or multiple-comparison correction was undertaken for the Figure 1B feature set. Panel C shows the direction and relative size of coefficients from the final standardised Logistic Regression model.

**Figure 1.**
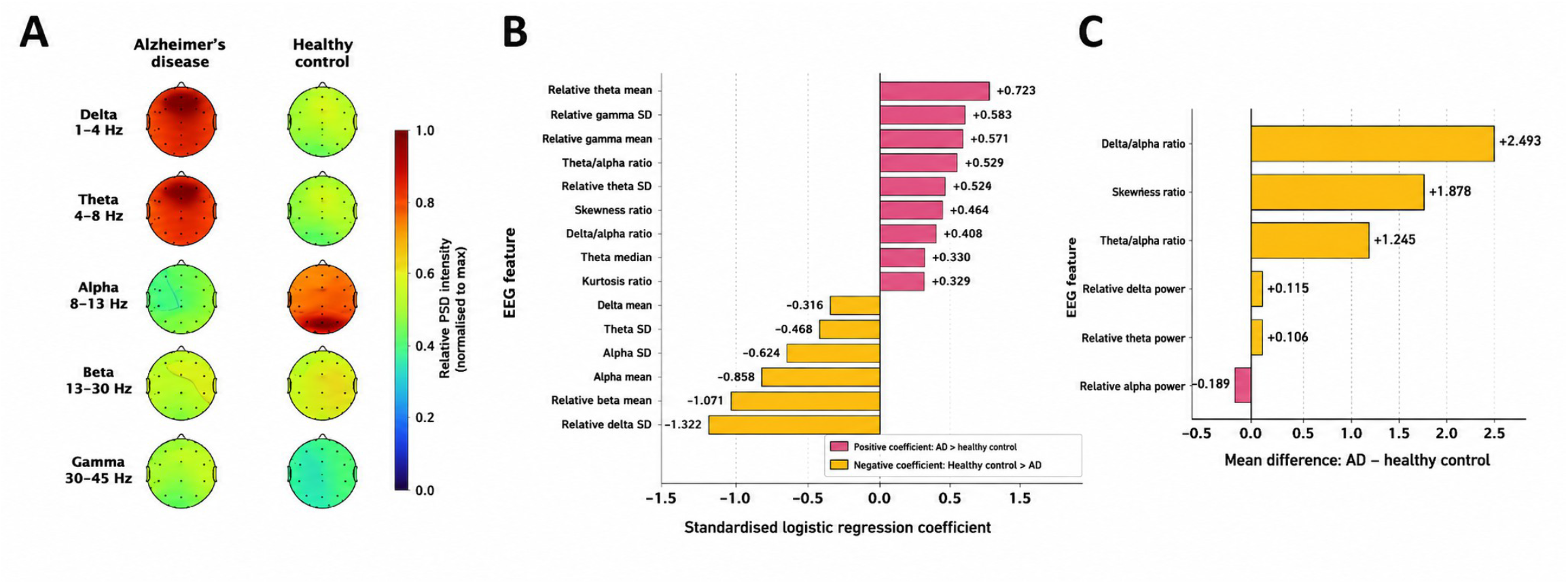
Spectral context and interpretable EEG feature behaviour. (A) Schematic adaptation of scalp power spectral density patterns described for the source dataset. Adapted from Miltiadous et al. [3]. Colours represent conceptual relative power spectral density intensity and do not show a new participant-level topographic re-analysis. (B) Descriptive mean differences between Alzheimer’s disease and healthy-control groups for selected native-scale EEG features. Positive values indicate higher mean feature values in the Alzheimer’s disease group. This panel is descriptive and does not represent formal between-group hypothesis testing. (C) Standardised Logistic Regression coefficients. Positive coefficients contribute toward the Alzheimer’s disease class in the fitted model, whereas negative coefficients contribute toward the healthy-control class.

### Model development and internal validation

All modelling was conducted in Python with scikit-learn [6]. The model comparison used five-fold stratified cross-validation. Stratification preserved the diagnostic class proportion across folds as far as possible, and the unit of splitting was the participant. This choice is important in EEG research. If overlapping segments from one participant are split across training and test data, a classifier can learn person-specific structure rather than disease-related structure. By using one row per participant, each held-out fold contained participants who were absent from the corresponding training set. The first EEG model was an Original Random Forest with 500 trees, maximum depth four, and balanced class weights. The second was a more constrained Random Forest with 300 trees, maximum depth two, minimum leaf size five, minimum split size ten, square-root feature sampling, and balanced class weights. The third model was L2-regularised Logistic Regression with a StandardScaler, balanced class weights, and a maximum of 1000 optimisation iterations. Logistic Regression was included because it provides an interpretable baseline. After feature standardisation, the coefficient sign indicates whether a feature shifts the fitted log-odds toward Alzheimer’s disease or toward healthy control, while the absolute coefficient size gives a rough indication of relative contribution within the fitted model. Two additional models were used as validation controls. The age-only baseline used participant age as its sole predictor. The EEG-plus-age Logistic Regression model appended age to the EEG feature matrix. These comparisons were designed to test a narrow but important possibility that diagnostic separation could arise primarily from age rather than from EEG-derived information. Accuracy, balanced accuracy, F1 score, and ROC-AUC were calculated for every fold. Mean test performance across folds is reported in Table 2. Training metrics were also inspected to identify overfitting, although they were not used as the primary evidence of performance. For the selected EEG Logistic Regression model, cross-validated predicted probabilities from held-out observations were pooled to create an out-of-fold ROC curve. Cross-validated class predictions were used to construct a pooled confusion matrix. This produces a useful visual summary of how the classifier behaved across all 65 participants, but it yields a different AUC summary from the mean of the five fold-specific AUC estimates. Both are reported in the manuscript for transparency rather than being treated as interchangeable.

**Table 1.** Dataset characteristics and EEG feature-engineering workflow for Alzheimer’s disease versus healthy-control classification.

| Component | Details |
| --- | --- |
| Source dataset | OpenNeuro ds004504, originally described by Miltiadous et al. |
| Full cohort | 88 participants: 36 Alzheimer's disease, 23 frontotemporal dementia, and 29 healthy controls. |
| Analytical cohort | 65 participants: 36 Alzheimer's disease and 29 healthy controls. |
| Recording condition | Resting-state, eyes-closed scalp EEG. |
| EEG montage | 19-channel international 10–20 scalp montage. |
| Sampling rate | 500 Hz. |
| EEG files used | Preprocessed EEGLAB derivative .set files. |
| Secondary standardisation | EEG-channel selection, average reference, and 1–45 Hz band-pass filtering. |
| Spectral estimation | Welch power spectral density estimation using 2-second segments. |
| Frequency bands | Delta 1–4 Hz; theta 4–8 Hz; alpha 8–13 Hz; beta 13–30 Hz; gamma 30–45 Hz. |
| Extracted features | Absolute bandpower mean, SD, median and maximum; relative bandpower mean and SD; theta/alpha ratio; delta/alpha ratio; slow/fast ratio; signal summary measures. |
| Final feature matrix | 65 participants × 37 EEG-derived predictor features. |
| Deployment output | Standardised Logistic Regression model deployed in a Gradio/Hugging Face research prototype. |

**Table 2.** Five-fold stratified cross-validation performance for Alzheimer’s disease versus healthy-control classification.

| Model | Feature set | Accuracy | Balanced accuracy | F1 score | ROC-AUC | Key interpretation |
| --- | --- | --- | --- | --- | --- | --- |
| Original Random Forest | EEG features | 0.754 | 0.757 | 0.757 | 0.851 | Strong training overfit; train accuracy 1.000. |
| Conservative Random Forest | EEG features | 0.785 | 0.791 | 0.780 | 0.875 | Reduced overfitting and improved test performance. |
| Logistic Regression | EEG features | **0.846** | **0.848** | **0.849** | **0.948** | Best overall EEG-only |
| Logistic Regression | Age only | 0.523 | 0.528 | 0.538 | 0.560 | Near-chance baseline. |
| Logistic Regression | EEG features + age | 0.846 | 0.848 | 0.849 | 0.943 | Age did not improve EEG-only classification. |

### Prototype development and technical testing

The final EEG Logistic Regression pipeline and feature-column list were saved and incorporated into a Gradio application [7]. The app offers a dataset-subject demonstration, feature CSV upload, and an experimental raw EEG upload route for .edf, .set, or .fif files. A second tab accepted a feature CSV. This was included for users who can reproduce the same feature schema externally. The app checks that required feature columns are present and reorders them to match the model’s training input. A third, experimental tab accepted EEG files in .edf, .set, or .fif format. The raw-file pathway attempted to load the recording, select EEG channels, apply the same secondary standardisation, extract the required feature set, and pass the resulting row to the saved classifier. The raw EEG route is experimental because the model was trained on eyes-closed, 19-channel, 500 Hz clinical EEG from one dataset. External recordings may differ in montage, sampling rate, reference, artefact profile, and task condition. An eyes-closed PhysioNet EDF file was used only as a technical smoke test of upload and processing, not as clinical validation [8].

## Results

### Spectral feature behaviour

Figure 1B presents descriptive mean differences for selected model-relevant features. Relative to healthy controls, the Alzheimer’s disease group showed higher mean delta/ alpha ratio (+2.493), slow/fast ratio (+1.828), theta/alpha ratio (+1.245), relative delta power (+0.115), and relative theta power (+0.106). Relative alpha power was lower in the Alzheimer’s disease group (−0.189). These values are presented in their native feature scales. The ratio features therefore appear numerically larger than relative-power differences and should not be compared as though they represent a common effect-size metric. No feature-level hypothesis testing or false-discovery-rate correction was performed because the primary purpose of this analysis was classifier development and model interpretation rather than confirmatory biomarker discovery. The observed directions are broadly aligned with the slowing pattern previously reported in Alzheimer’s disease EEG literature and in the ds004504 dataset paper [3]. However, the present descriptive comparisons should not be interpreted as proof that individual features differed significantly between diagnostic groups in this sample.

### Model comparison

The Original Random Forest produced a mean test accuracy of 0.754, balanced accuracy of 0.757, F1 score of 0.757, and ROC-AUC of 0.851. Its training accuracy and training ROC-AUC were both 1.000, indicating that the initial forest was memorising much of the training structure. The Conservative Random Forest reduced this gap. Its mean test accuracy increased to 0.785, balanced accuracy to 0.791, F1 score to 0.780, and ROC-AUC to 0.875, while training accuracy fell to 0.888. This was a useful improvement, but the tree-based models remained below the regularised linear model. EEG Logistic Regression was the strongest model in the final comparison. Mean test accuracy was 0.846, balanced accuracy 0.848, F1 score 0.849, and mean fold-wise ROC-AUC 0.948. The model retained a training-test difference, with training accuracy of 0.954 and training ROC-AUC of 0.986, so the result should not be read as evidence of perfect generalisation. Still, the lower apparent complexity of the model and the agreement between predictive performance and interpretable feature directions made it the most suitable pipeline for deployment. The age-only baseline performed close to chance. It produced a mean test accuracy of 0.523, balanced accuracy of 0.528, F1 score of 0.538, and ROC-AUC of 0.560. When age was appended to the EEG features, accuracy, balanced accuracy, and F1 score remained identical to the EEG-only Logistic Regression model, while mean ROC-AUC was slightly lower at 0.943. The practical conclusion is limited but clear. In this cohort, adding age did not improve the model, and age alone could not reproduce the discrimination obtained from EEG features. Figure 2 summarises the performance evidence in three ways. Panel A displays the pooled out-of-fold ROC curve for EEG Logistic Regression, with a pooled AUC of 0.916. Panel B shows that 25 of 29 healthy-control recordings and 30 of 36 Alzheimer’s disease recordings were classified correctly. This gives specificity of 86.2% and sensitivity of 83.3%. Panel C compares mean fold-wise ROC-AUC values. The highest bar is EEG Logistic Regression (0.948), followed closely by EEG plus age (0.943), with age-only classification near the 0.50 chance line.

**Figure 2.**
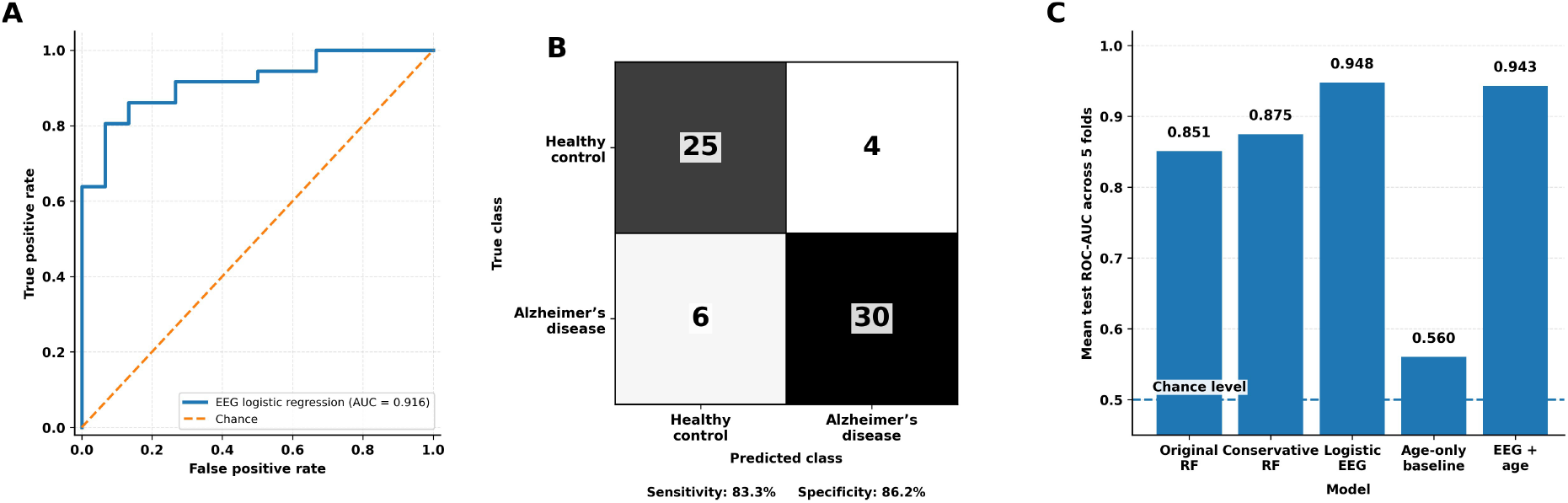
Model performance under five-fold stratified cross-validation. (A) Pooled out-of-fold ROC curve for EEG Logistic Regression. (B) Pooled cross-validated confusion matrix. (C) Mean fold-specific ROC-AUC values for the compared models. The dashed line marks a chance-level ROC-AUC of 0.50.

### Model interpretation and public deployment

Coefficient inspection added context to the final prediction model. Positive coefficients included relative theta mean (+0.723), relative gamma standard deviation (+0.583), relative gamma mean (+0.571), theta/alpha ratio (+0.529), relative theta standard deviation (+0.464), slow/fast ratio (+0.445), and delta/alpha ratio (+0.409). Several alpha-related features carried negative coefficients, including alpha maximum (−0.836) and alpha standard deviation (−0.624). Relative beta mean (−1.071) and relative delta standard deviation (−1.322) were also negative. These values should be read as properties of one multivariable model after feature standardisation, not as independent disease effects. The coefficient pattern provides model-level interpretive context rather than independent feature-level evidence of diagnostic group differences. Relative theta and slow-to-alpha ratios contributed positively toward the Alzheimer’s disease class, while stronger alpha-related features tended to support the healthy-control class in the fitted Logistic Regression model. These coefficient directions are consistent with the broad spectral slowing pattern described in prior Alzheimer’s disease EEG literature.

However, coefficients reflect the behaviour of correlated features within one multivariable classification model and should not be interpreted as independent univariate disease effects. Gamma-derived coefficients are reported for transparency but require particular caution because high-frequency scalp EEG can be influenced by residual muscle activity and recording artefact. The final app was successfully launched locally in Kaggle and then deployed as a public Hugging Face Space. It allows a user to select a source-dataset participant, upload a compatible feature CSV, or test the experimental raw-EEG path. The app is available at https://huggingface.co/spaces/Atrim777/Screening. Its public availability is an extension of the research workflow, not evidence that it can be used outside the training context. The interface states that it should not be used to diagnose Alzheimer’s disease.

## Discussion

This study took an open clinical EEG dataset through a complete small-scale machine-learning workflow: reproducible loading, secondary standardisation, subject-level feature engineering, model comparison, validation controls, interpretation, and public deployment. Its contribution lies in bringing these elements together. Rather than ending with an accuracy score in a coding environment, the work produced an interface that makes the model and its limits easier to inspect. That does not make the app clinically ready, but it makes the analytical choices more visible. Miltiadous et al. [3] provided the source dataset and an important benchmark, but this study does not claim an exact replication of their approach. Their analysis used relative bandpower, four-second epochs with 50% overlap, and leave-one-subject-out validation. Here, each participant contributed one feature row, and models were evaluated with five-fold stratified cross-validation. The question was therefore different: can a compact, participant-level feature representation support an interpretable AD versus healthy-control classifier within the same dataset? The participant-level design was important because EEG segments from the same recording are highly correlated. Splitting such segments between training and test data can inflate performance by allowing a classifier to learn person-specific patterns. Using one row per participant reduces that risk, although it leaves a modest effective sample size and keeps the study firmly in proof-of-concept territory. The model comparisons also sharpened interpretation. The original Random Forest achieved reasonable held-out performance but perfect training metrics, which suggested overfitting. Restricting tree depth and leaf size reduced this gap. Logistic Regression then performed best on the reported test metrics while remaining easier to interpret. The age-only baseline was particularly useful. Age is a plausible confound in an Alzheimer’s disease dataset, yet it performed near chance here, and adding age did not improve the EEG-only model. This does not rule out all confounding, but it weakens a simple age-based explanation for the observed classification signal. Interpretability was treated as part of the result. The final model assigned weight to theta-related, alpha-related, and slow-to-fast ratio features whose descriptive directions were broadly compatible with established reports of EEG slowing in Alzheimer’s disease. These findings should not be read as confirmed feature-level biomarkers. Figure 1B presents descriptive mean differences without univariate hypothesis testing, effect-size estimates, or multiple-comparison correction. It was included to make the feature space transparent and to contextualise the fitted model, not to establish statistically significant group differences. The public prototype adds a practical dimension. The feature-CSV route is relatively reproducible because it expects the same feature schema used during training. The raw EEG upload route is more experimental. A file can load successfully while still being unsuitable for interpretation because of differences in montage, sampling frequency, reference, task condition, artefact profile, or clinical population. Several limitations remain. The binary cohort contained 65 participants from one source context, with no external cohort or prospective clinical evaluation. Model selection and performance estimation relied on the same internal cross-validation framework; nested cross-validation or an independent development set would provide a stricter estimate. Finally, the application must not be mistaken for a diagnostic product. Alzheimer’s disease diagnosis requires symptoms, longitudinal history, cognitive assessment, clinical examination, and, where appropriate, biomarker or imaging evidence. The app therefore refers to AD-like and healthy-control-like EEG profiles rather than diagnostic verdicts.

Future work would require external validation, calibration analysis, uncertainty estimates, prospective testing, privacy safeguards, and a clearly defined clinical workflow.

## Conclusion

This project demonstrates a reproducible route from open resting-state EEG data to an interpretable public research prototype. Using the Alzheimer’s disease and healthy-control subset of OpenNeuro ds004504, it extracted subject-level spectral and oscillatory features, compared model families, tested age as a competing explanation, and selected a regularised Logistic Regression classifier for deployment. The EEG-only model achieved the strongest mean cross-validated performance, while age alone was near chance and did not improve the EEG model. The fitted feature directions were consistent with a slow-wave shift and reduced alpha-related activity in Alzheimer’s disease. The work is a transparent proof of concept for research and education, not a diagnostic tool.

## Data and prototype availability

The de-identified source data are publicly available through OpenNeuro as ds004504 [3]. The deployed research prototype is available at [https://huggingface.co/spaces/Atrim777/Screening]. The application is provided for educational and research demonstration only. It must not be used as a standalone clinical diagnostic or screening tool.

